# Mutational inactivation of Apc in the intestinal epithelia compromises cellular organisation

**DOI:** 10.1101/2020.06.10.143800

**Authors:** Helena Rannikmae, Samantha Peel, Simon Barry, Inderpreet Sur, Jussi Taipale, Takao Senda, Marc de la Roche

## Abstract

The tumour suppressor adenomatous polyposis coli (Apc) regulates diverse effector pathways essential for cellular homeostasis. Truncating mutations in Apc, leading to the loss of its Wnt pathway and microtubule regulatory domains, are oncogenic in human and murine intestinal epithelia and drive malignant transformation. Whereas uncontrolled proliferation via Wnt pathway deregulation is an unequivocal consequence of oncogenic Apc mutations, it is not known whether loss of its other control systems contribute to tumorigenesis. Here we employ *in vitro* models of tumorigenesis to unmask the molecular barriers erected by Apc that maintain normal epithelial homeostasis in the murine intestinal epithelia. We determine that (i) enterocyte proliferation, (ii) microtubule dynamics and (iii) epithelial morphology are controlled by three independent molecular pathways, each corrupted by oncogenic Apc mutations. The key result of the study is to establish that Apc regulates three individual biological fates in the intestinal epithelia, through three distinct effector pathways, a significant advance to our understanding of normal tissue homeostasis, the molecular architecture of epithelial tissue and the aetiology of intestinal cancer.

## Introduction

The intestinal tract (small intestine and colon) hosts a highly dynamic enterocyte monolayer that undergoes complete self-renewal every 3-5 days. The basic units of the intestinal epithelium are adjacent invaginations, termed crypts of Lieberkühn (Fig. 1A, B), each of which serves as a semi-autonomous cell production factory with a remarkably high proliferation rate - along the murine intestinal tract, crypts are composed of an average of 700 cells that produce up to 20 cells per hour in the small intestine or 7 cells per hour in the colon (de Rodriguez et al., 1978; Potten et al., 1982; Sunter et al., 1979). Throughout the enterocyte monolayer, each cell is spatially restricted, selective for homo- and hetero-typic cell-specific interactions, and is highly polarised with defined apical, lateral and basal faces, all characteristics critical to epithelial barrier and transport functions. The hierarchal organisation of the enterocyte monolayer, from the stem cells at the base of crypts to differentiated cells within the gut lumen, is achieved through molecular control of the balance between rapid cellular proliferation and morphological organisation of the epithelial monolayer (Gehart and Clevers, 2019).

**Figure 1.**
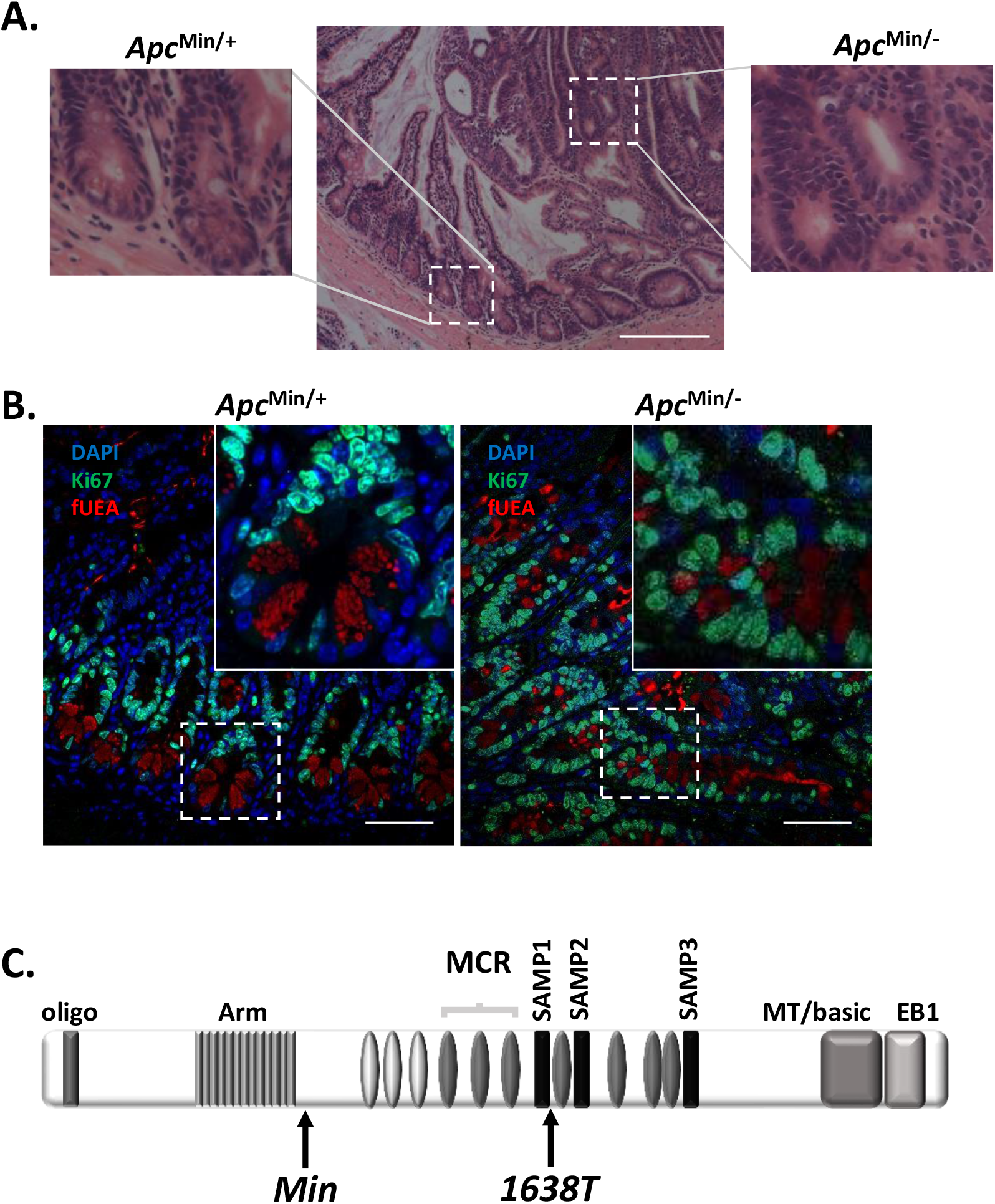
Compromised morphology of the monolayer and cellular organisation in Apc deficient intestinal epithelia. **A.** Haematoxylin and eosin stain of normal *Apc*^*Min*/+^ murine intestinal epithelia and adjacent *Apc*^Min/−^ tumour. Outsets illustrate crypt units for normal *Apc*^*Min*/+^ murine intestinal epithelia and gland-like structures in *Apc*^*Min*/−^ tumours. Scale bars, 200 μm. **B.** Fluorescent confocal microscope imaging of small intestinal epithelial sections from an *Apc*^Min/+^ mouse. *Left panel* - normal, haplo-sufficient *Apc*^Min/+^ tissue. *Right panel* - *Apc*^Min/−^ tumour. Paneth cell vesicles are marked with fUEA (red), fluorescent antibody to Ki67 (green) marks cycling cells and DAPI (blue) labels nuclei. Scale bars, 50 μm. **C.** Domain structure of Apc showing protein interaction domains labelled as: oligo - oligomerisation domain; Arm – armadillo repeat domain; Axin-binding SAMP domain 1-3; MT/basic – the microtubule binding domain containing basic amino acids; EB1 – EB1-binding domain; ovals refer to 15- and 20-amino acid β-catenin binding domains (grey and dark grey, respectively). MCR is the position of the corresponding <u>m</u>utational <u>c</u>luster <u>r</u>egion in human APC. Also shown are the relative positions of the germline *Min* and *1638T* mutations in Apc.

Malignant transformation as a result of mutational inactivation of the tumour suppressor gene *adenomatous polyposis coli* (*APC*) compromises tissue organisation of the intestinal epithelium (Dow et al., 2015; Kinzler, Kenneth W. and Vogelstein, 1996; Volgestein and Fearon, 1990). Somatic mutations in *APC* are widely regarded as the initiating event of 80-90 % of sporadic colon cancers (Groden et al., 1991). Perhaps surprisingly, mutational inactivation of APC reveals an oncogenic vulnerability largely restricted to the intestinal epithelium. Thus, individuals with familial adenomatous polyposis (FAP) that are heterozygous for a germline mutation inactivating one allele of *APC* (Su et al., 1992) exhibit spontaneous loss of heterozygosity that leads to hundreds of tumours, all of which are restricted to the intestinal epithelium. The murine model of FAP, *Apc*^Min/+^ (multiple intestinal neoplasia; *Min*), follows a similar pattern of tumour development—despite mono-allelic inactivation of *Apc* in every cell in the body, tumorigenesis is almost exclusive to the intestinal epithelium (Moser et al., 1990; Moser et al., 1995; Su et al., 1992).

Apc is a large multi-domain protein that governs a plethora of effector pathways regulating cellular and tissue homeostasis (Nelson and Näthke, 2013). Apc’s molecular roles are generally ascribed to the regulation of Wnt pathway activity, a key determinant of stem cell multipotency and proliferation within the crypt. Pathway activity is sustained within the stem cell niche by redundant sources of Wnt ligands derived from adjacent Paneth cells or the underlying mesenchyme (Aoki et al., 2016; Farin et al., 2012; Gregorieff et al., 2005; Stzepourginski et al., 2017; Valenta et al., 2016; Zou et al., 2018) and potentiated by cellular engagement of R-spondins, also derived from specific mesenchymal cells (Yan et al., 2017).

APC inactivation is one of the earliest known lesions in colorectal cancer and follows an unusual pattern of somatic changes – at least one APC allele harbours mutations that are largely confined to a short segment within *exon 15* of the gene referred to as the mutation cluster region (MCR; Fig. 1C, S1), resulting in the expression of truncated Apc. The other allele is most often silenced or incurs the same or a more severe truncating mutations (Crabtree et al., 2003; Lamlum et al., 1999; Rowan et al., 2000). Truncating mutations in *exon 14* of the mouse *Apc* gene found in the *Min* mouse line (Fig. 1C, S1), equivalent to human *exon 15*, display many features common with human colorectal cancer.

The truncated Apc protein lacks regulatory protein-protein interaction domains for the Wnt pathway regulators β-catenin and Axin (Fig. 1C, S1), explaining oncogenic Wnt pathway activation upon loss of heterozygosity in murine models. Extensive investigation of oncogenic Wnt pathway activity in cells lacking APC has ascribed a key role in the regulation of intestinal epithelial cell proliferation through the Wnt pathway target gene c-*Myc* (Dave et al., 2017; He et al., 1998; Oskarsson and Trumpp, 2005; Sansom et al., 2007; Sur et al., 2012).

Truncated Apc protein also lacks the C-terminal microtubule <u>e</u>nd <u>b</u>inding protein 1 (EB1) binding domain and a basic domain thought to bind directly to microtubules (Fig. 1C) (Deka et al., 1998). However, the molecular consequence of C-terminal Apc truncations and removal of the microtubule and EB1 binding domains is controversial. Apc mediated stabilisation of microtubules via its C-terminal domains supports the establishment of parallel arrays of microtubules in a polarised cell (Mogensen et al., 2002; Zumbrunn et al., 2001) and APC is known to regulate cytoskeletal rearrangements that accompany cell motility, cell division and tissue organisation through control of microtubule dynamics (Moseley et al., 2007; Munemitsu et al., 1994; Näthke, 1996; Smith et al., 1994). It is not clear, if the truncating mutations in APC decrease its binding to microtubules or its capacity to stabilize the microtubule ends (Munemitsu et al., 1994; Smith et al., 1994)(Karin Kroboth et al., 2007; Zumbrunn et al., 2001). Furthermore, loss of Apc C-terminal microtubule and EB1 interaction domains in mouse embryonic fibroblasts and differentiated embryonic stem cells does not affect the distribution of β-tubulin, EB1 and Apc (Lewis et al., 2012; Smits et al., 1999).

*In vivo* mouse models have investigated whether loss of Apc’s C-terminal microtubule and EB1 binding domains are sufficient to drive intestinal epithelial tumorigenesis. *Apc*^1638T/1638T^ mice express a version of Apc lacking the C-terminal domains but retaining the ability to regulate Wnt pathway activity (Fig. 1C, S1) and do not present with intestinal epithelial tumours (Smits et al., 1999). Conversely, *Apc*^ΔSAMP/+^ mice expressing a version of Apc unable to regulate Wnt pathway activity but retaining the microtubule and EB1 binding domains (Fig. S1) develop tumours with the same frequency and kinetics as the corresponding *Apc*^1322/+^ mice that express Apc lacking these domains (Fig. S1) (Lewis et al., 2012). Thus, Apc’s ability to interact with microtubules and EB1 does not, on its own, drive intestinal epithelial tumorigenesis. Nonetheless, the potential contribution of loss of the APC microtubule and EB1 binding domains to intestinal tumorigenesis has not been determined.

Herein, we stratify functions of Apc in the murine intestinal epithelia by defining the molecular and phenotypic differences in the small intestinal epithelia and corresponding organoids that arise upon loss-of-function. In addition to deregulated cell proliferation, we find that Apc inactivation disrupts intestinal epithelial morphology and compromises microtubule dynamics in component enterocytes. Although the three emergent malignant properties are the direct consequence of Apc inactivation, they are controlled by different molecular systems. Therefore, (i) enterocyte proliferation, (ii) microtubule dynamics and (iii) epithelial morphology are regulated by three separate effector pathways, under the control of Apc, that bulwark normal intestinal epithelial homeostasis against malignant transformation.

## Results

### Compromised intracellular organisation and tissue morphology in *Apc*^Min/−^ tumours

Over the course of 110 days, *Apc*^Min/+^ mice develop 30-40 adenomas in the small intestine, the result of loss of heterozygosity of the wild type *Apc* allele (Moser et al., 1995; Su et al., 1992). Such *Apc*^Min/−^ tumours are composed of gland-like structures that maintain a columnar epithelial monolayer yet lack the morphological hallmarks of crypt and villus compartments and the hierarchal cellular organisation of the wild type epithelia (Fig. 1A). For instance, Ki67^+^ proliferative stem cells and the transit amplifying cellular compartment, normally disposed supra-basally within crypts, are instead interspersed throughout the monolayer of the tumour gland-like structures (Fig. 1B). Moreover, a fluorescent probe of apically-localised secretory vesicles (fluorescently-labelled *Ulex Europaeus* agglutinin; fUEA) found in the mechanically rigid, keystone-shaped Paneth cells in wild-type tissues (Langlands et al., 2016), indicates that these cells are interspersed throughout the *Apc*^Min/−^ glandular monolayer, are of variable shapes and fail to maintain apical vesicle localisation (Fig. 1B). We also note that, as opposed to wild type enterocytes, cells within the *Apc*^Min/−^ tumour contain nuclei of variable shapes and sizes that do not align along the plane of the monolayer. We conclude that, in addition to driving de-regulated epithelial cell proliferation and tissue morphology, Apc inactivation compromises molecular barriers maintaining some aspects of intracellular organisation.

### Defective regulation of microtubule function in *Apc*^Min/−^ tumours

The cytoskeleton provides the physical framework for intracellular organisation and cell polarity defined by dynamic polymerisation/depolymerisation of actin and tubulin monomers (Li and Gundersen, 2008; Rodriguez-Boulan and Macara, 2014). Apc harbours an array of protein-protein interaction domains with established roles in regulating F-actin and microtubule dynamics within intestinal epithelial cells (Fig. 1C)(Kawasaki et al., 2000; Munemitsu et al., 1994; Näthke, 2004; Rosin-Arbesfeld et al., 2001; Tirnauer, 2004; Zumbrunn et al., 2001). We examined the localisation of the cytoskeleton in intestinal epithelial and *Apc*^*Min*/−^ tumour cells, using a series of fluorescent probes for F-actin, microtubules and known protein interactors. Consistent with a previous study (Fatehullah et al., 2013), *Apc*^Min/−^ tumour cells maintained the correct disposition and configuration of actin cytoskeletal components— F-actin was concentrated along the apical face of the epithelial cells (Pelaseyed and Bretscher, 2018), the tight junction organiser ZO-1 was positioned apically at cellular junctions (Lee et al., 2018) and integrin-β4, which anchors enterocytes to the underlying lamina propria, localised to the cell base (Fatehullah et al., 2013) (Fig. 2A).

**Figure 2.**
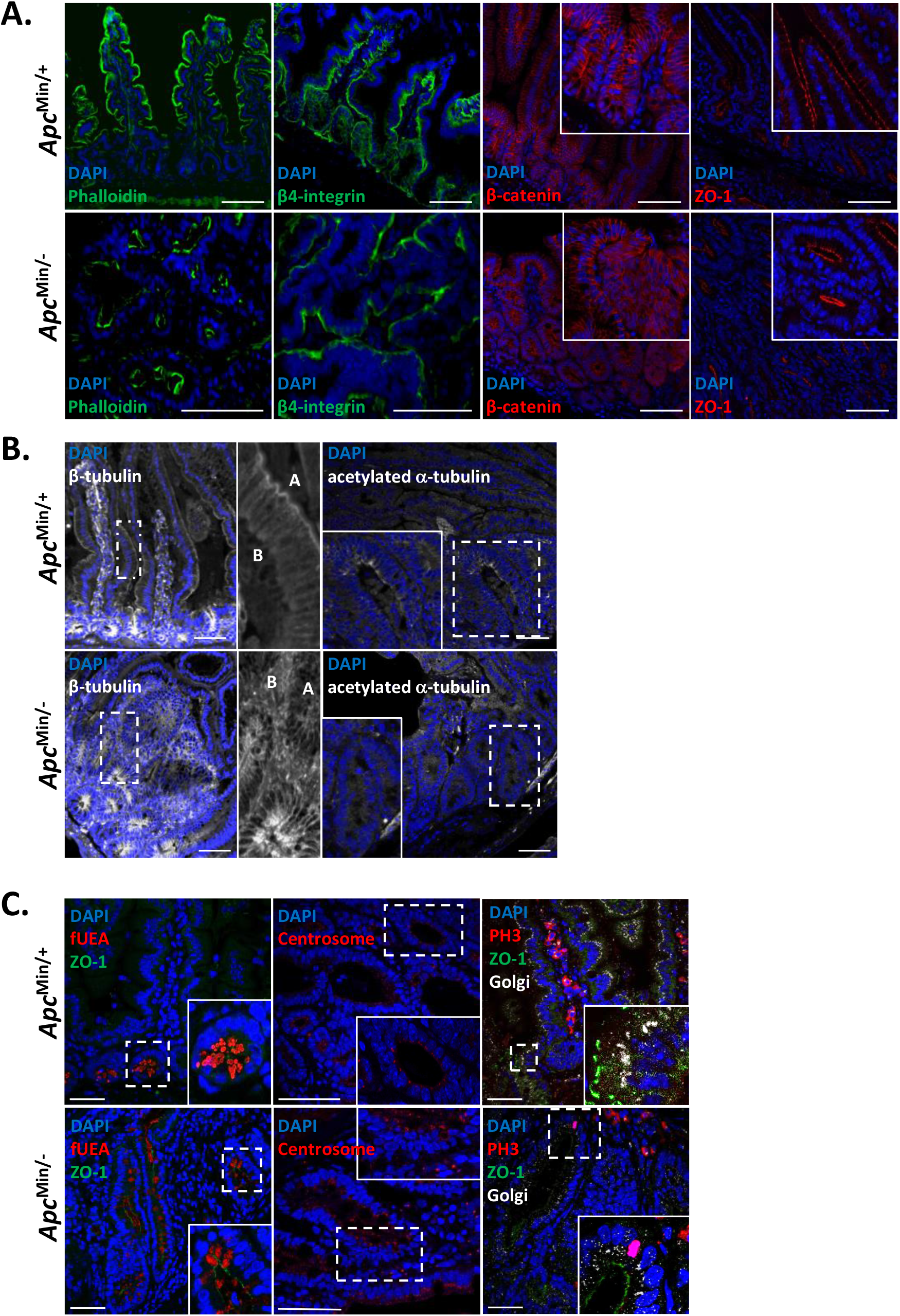
Apc inactivation does not affect the localisation of the actin cytoskeleton yet compromises microtubule dynamics. Fluorescence confocal microscopy of small intestinal epithelia sections (from *Apc*^Min/+^ mouse; top panels) and *Apc*^Min/−^ tumours, bottom panels. **A.** Leftmost sections were labelled with fluorescent phalloidin (*green*); the subsequent pairs of sections were labelled with antibodies to β4-integrin (green), β-catenin (red) and ZO-1 (red). **B.** Leftmost section was labelled with an antibody to β-tubulin. Expanded view to the right, “A” marks the apical domain of cells and “B”, the basal domain. Rightmost section was labelled with an antibody to acetylated α-tubulin. **C.** Left panels – sections labelled with fUEA (red) and an antibody to ZO-1 (green); middle panels – sections labelled for the centrosome using an antibody to pericentrin (red); right panels – sections labelled with antibodies to the Golgi resident protein ZFPL1 (red), the cell division marker PH3 (white) and ZO-1 (green). All sections were co-labelled with DAPI. Scale bars, 100 μm.

In contrast, components of the microtubule cytoskeleton in *Apc*^*Min*/−^ tumour cells were disorganised; microtubules, normally orientated along the apical-basal axis were instead disjointed and diffuse (Fig. 2B). We used an antibody raised against the acetylated form of α-tubulin and found that the signal was concentrated at the apical domain of cells, in line with previously published data (Quinones et al., 2011). However, in tumour cells, acetylated α-tubulin was instead de-localised and diffuse (Fig. 2B). We also determined the localisation of intracellular organelles whose location and disposition are dependent on the microtubules. Predictably, we found that the normally strict basal positioning of nuclei, apical positioning of intracellular vesicles and the supra-apical localisation of the Golgi resident protein ZFLP1 and the centrosome marker pericentrin in wild type intestinal epithelia was lost in *Apc*^*Min*/−^ tumour cells (Fig. 2C). To preclude de-localisation of the Golgi as the consequence of cells undergoing cell division, we co-stained intestinal epithelial sections with an antibody to the mitotic marker phospho-histone 3 (PH3; Fig. 2C) - tumour cells that displayed de-localised Golgi did not express detectable levels of PH3. Taken together, our data supports normal localisation of the actin cytoskeleton and associated components within *Apc*^*Min*/−^ intestinal epithelial tumour cells, whereas the localisation and functional integrity of the microtubules is compromised.

We reasoned that the C-terminal microtubule and EB1 binding domains of Apc may be critical for the regulation of the microtubule cytoskeleton. The *Apc*^1638T/1638T^ mouse strain is homozygous for a truncating mutation in Apc that deletes the C-terminal microtubule and EB1 binding domains. However, Apc^1638T^ protein retains the Axin interaction domain unlike that expressed in *Apc*^*Min*/+^ mice (Fig. 1C) and therefore retains regulatory control over Wnt pathway activity; as a result, *Apc*^1638T/1638T^ mice do not develop intestinal epithelial tumours (Smits et al., 1999). Since the small intestine epithelia of *Apc*^1638T/1638T^ mice exhibit normal localisation of intact Golgi and fUEA-positive Paneth cell vesicles (Fig. S2), we conclude that loss of the Apc microtubule and EB1 binding domains alone does not compromise regulation of the microtubule cytoskeleton or intestinal epithelial morphology.

### Organoids accurately recapitulate the molecular and phenotypic consequences of APC inactivation in the intestinal epithelium

We generated organoid lines from wild-type, *Apc*^Min/+^ intestinal epithelia and *Apc*^Min/−^ tumour cells as an experimentally tractable model system for determining the molecular mechanisms linking Apc to microtubule integrity and epithelial morphology. Organoids derived from normal tissue form an epithelial monolayer, replete with crypts, that maintains the three-dimensional cellular organisation and hierarchy found *in vivo*. In contrast, tumouroids, organoids derived from *Apc*^Min/−^ tumour cells, form cystic structures lacking morphological features of the intestinal epithelial monolayer such as crypts (Sato et al., 2011).

Using a series of fluorescent probes, we found that F-actin and associated molecular components, ZO-1 and integrin-β4, maintained their intracellular localisation in both organoid and tumouroid cells (Fig. 3A). However, consistent with our observations in intestinal epithelial tissue from *Apc*^Min/−^ tumours, the organisation and function of the microtubule cytoskeleton was compromised. Notably, β-tubulin was no longer polarised in microtubules along the apical-basal axis of cells but was instead dispersed throughout all of the cells, acetylated α-tubulin was de-localised (Fig. 3B), nuclei varied in shape and size and did not follow the plane of the tumouroid monolayer and centrosomes and Golgi were split into multiple puncta that were distributed throughout the cell body (Fig. 3C); within individual tumouroid cells, the Golgi and the centrosome were no longer positioned apically in over 40% of cases (Fig. 3D).

**Figure 3.**
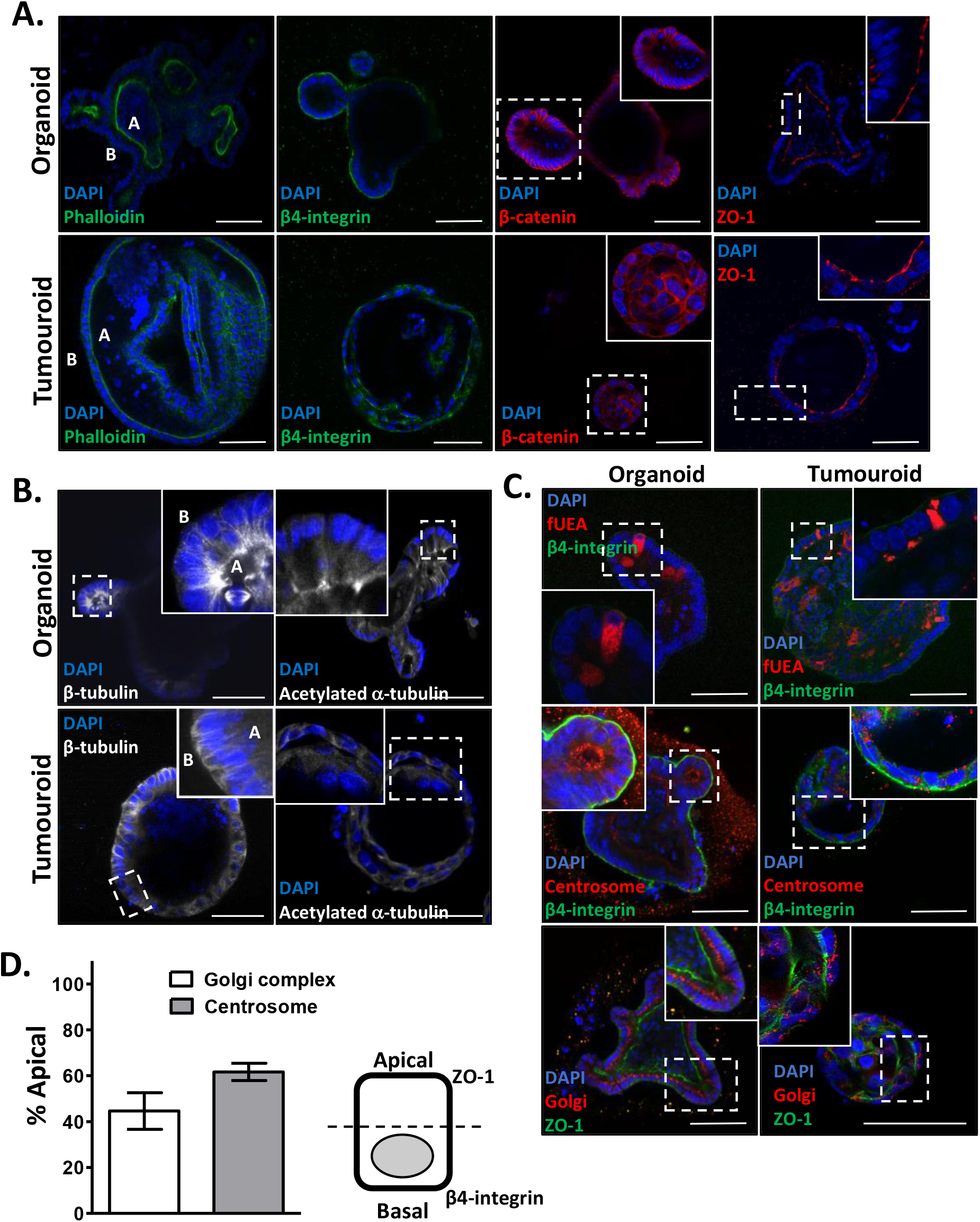
Organoids recapitulate the consequences of Apc inactivation in the intestinal epithelia. **A.** Fluorescence confocal microscopy of small intestinal epithelial organoids (top panels) and *Apc*^Min/−^ tumouroids (bottom panels). Cells were labelled with fluorescent phalloidin (green) and antibodies to β4-integrin (green), β-catenin (red) and ZO-1 (red) as marked. On the left panel “A” marks the apical domain of cells and “B”, the basal domain. **B.** Immunofluorescence using antibodies to β-tubulin and acetyl-tubulin as marked. “A” marks the apical domain of cells and “B”, the basal domain. **C.** Fluorescent sections of small intestinal epithelial organoids (*left panels*) and *Apc*^Min/−^ tumouroids (*right panels*). Top panels were labelled with fUEA (red) and an antibody to β4-integrin (green); middle panels were labelled with antibodies to pericentrin (red) and β4-integrin (green); bottom panels were labelled with antibodies to ZFPL1 (red) and ZO-1 (green). All fluorescent sections were co-labelled with DAPI. Scale bars, 50 μm. **D.** The positioning of the Golgi complex and the centrosome was scored as apical or otherwise according to the scheme in the right panel for >200 cells from at least three independent experiments. Error bars ± SD.

Taken together, our organoid data confirms that Apc inactivation in the intestinal epithelial monolayer leads to deregulation of microtubule dynamics and loss of intracellular organisation with the absence of detectable effects on the actin cytoskeleton.

### Apc deficiency directly compromises intracellular organisation and tissue morphology

It is possible that intestinal epithelial tumours from 110-day old *Apc*^Min/+^ mice, and organoids derived from them, have acquired additional somatic changes that contribute to phenotype. To determine the immediate and direct effects of Apc inactivation we created a switchable organoid model of tumorigenesis that relies on the inducible expression of a previously-validated shRNA targeting Apc (Dow et al., 2015) (Fig. S3A). shApc expression in organoids depletes Apc expression concurrent with the expression of mCherry and leads to the intra-conversion of organoids into a cystic tumoroid structure (Fig. S3B-D). Importantly, we observe increased expression of the Wnt pathway target gene *c-Myc* (Fig. S3E).

Consistent with the appearance of *Apc*^Min/−^ tumours and tumoroids, Apc depletion in organoids resulted in mis-localisation of Paneth cell vesicles and Golgi and centrosome fragmentation and dispersion (Fig. 4A). Importantly, all hallmarks of intracellular disorganisation and compromised tissue morphology were reversed upon Apc re-expression, leading to the appearance of ‘normal’ organoids (Fig. 4A). Our switchable *in vitro* tumorigenesis model confirms that compromised epithelial morphology and intracellular disorganisation are a direct consequence of Apc inactivation.

**Figure 4.**
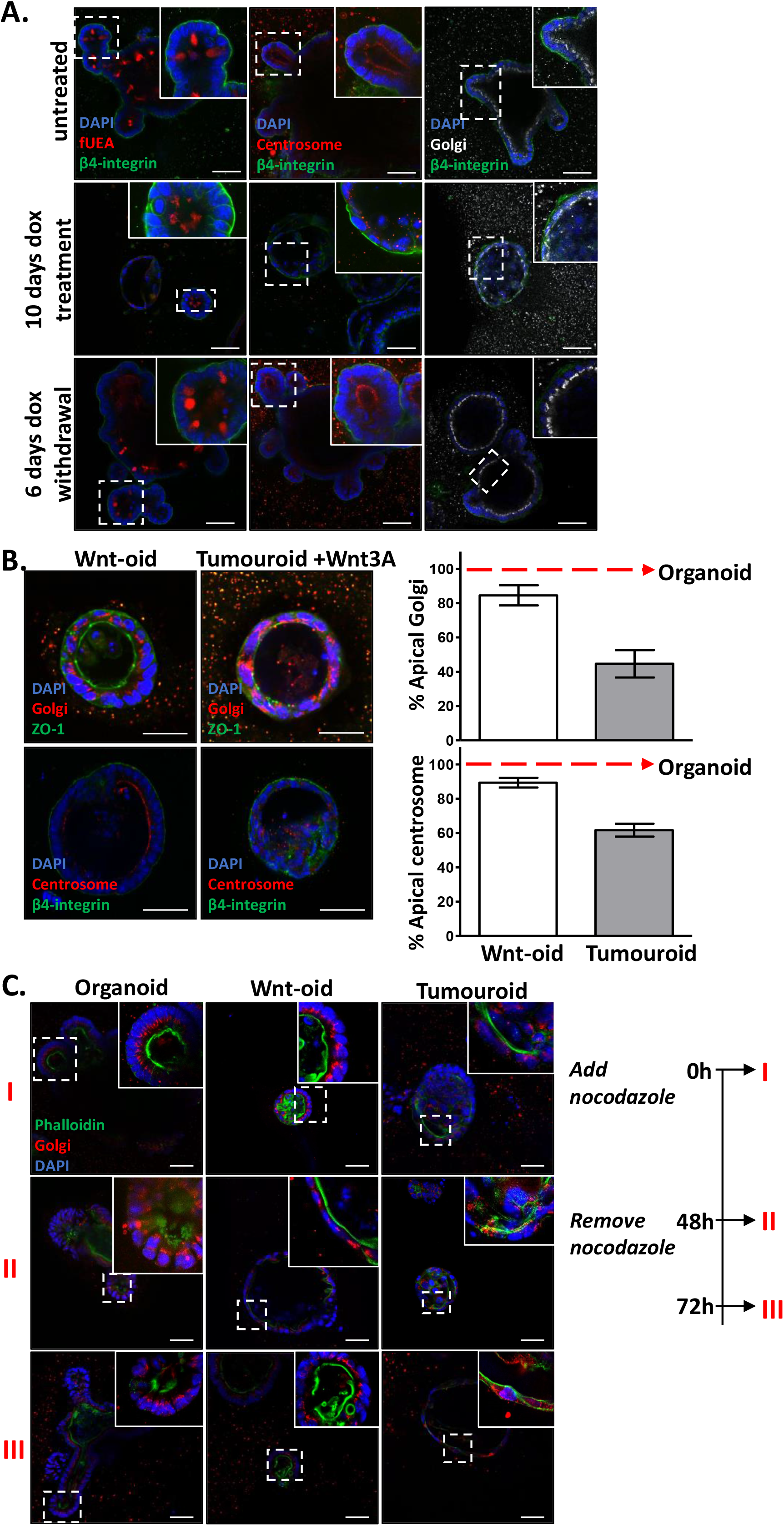
Switchable *in vitro* model of tumorigenesis recapitulates the consequences of Apc inactivation in the intestinal epithelia. **A.** An organoid line bearing pB-shApc, the *tet-on* inducible transgene system for induction of shApc expression untreated (top panels), treatment with doxycycline for 10 days (middle panels) or the former followed by doxycycline withdrawal for an additional 6 days (lower panels). *Left panels* – fluorescence confocal microscopy of organoids labelled with fUEA (red) and an antibody to β4-integrin (green); *middle panels* – organoids labelled with antibodies to pericentrin (red) and β4-integrin (green); *right panels* – organoids labelled with antibodies to ZFPL1 (white) β4-integrin (green). **B.** Fluorescence confocal microscopy of organoids and tumoroids treated with Wnt3A conditioned media for 72 hours. Right panels – quantification of apical localisation of centrosome and Golgi for the Wnt-oids and tumouroids treated with Wnt3A. Greater than 200 cells from three independent fluorescent sections were analysed using criteria described in Figure 3D. Error bars ± SD. **C.** Fluorescence confocal microscopy of organoids, Wnt-oids and tumoroids treated with nocodazole followed by fixation and/or withdrawal of drug as depicted in schematic. Images are representative of the behaviour of 50 organoids that were analysed per condition from at least two independent experiments. All sections were labelled with fluorescent phalloidin (*green*) and an antibody to ZFPL1 (red). All sections were co-labelled with DAPI. Scale bars, 50 μm.

### Apc regulation of intestinal epithelial morphology and microtubule dynamics are discrete

Ubiquitous activation of Wnt pathway activity in organoid cells by treatment with Wnt3A conditioned media leads to the intra-conversion of organoids into cystic tumouroid-like structures (Farin et al., 2012) that we refer to as Wnt-oids (Fig. 4B). Although the morphology of the Wnt-oid epithelial monolayer is compromised, they are distinct from tumouroids in that the Golgi, centrosome and Paneth cell vesicles retain their normal apical position in component cells (Fig. 4B) – greater than 80% of Wnt-oid cells show apical localisation of the Golgi and centrosome as opposed to less than 60% in tumouroid cells (Fig. 4B). We conclude that Apc regulation of intestinal epithelial morphology through Wnt pathway regulation is not coupled to its function in regulating microtubule dynamics and intracellular organisation.

We carried out the complimentary experiment, selectively deregulating microtubule dynamics in organoids and determining the consequence on epithelial morphology. We treated organoids with a low concentration (100 nM) of the microtubule depolymerising agent nocodazole (Vasquez et al., 1997) for 48 hours, a timepoint sufficient for the conversion of organoids to Wnt-oids with Wnt3A treatment. Treated organoid and Wnt-oid cells displayed the characteristic mis-localisation of fragmented Golgi and centrosomes that was reversed after 24 hours post-nocodazole withdrawal (Fig. 4C). Importantly, throughout the experiments, nocodazole-treated organoids maintain intestinal epithelial crypts structures (Fig. 4C) indicating that maintenance of intestinal organisation and microtubule dynamics are not dependent on one another. Combined with our Apc loss-of-function studies, these data suggest that Apc-dependent control of intracellular organisation and epithelial morphology rely on independent molecular circuits.

### Apc control of intestinal epithelial morphology is independent of the Wnt pathway target gene c-Myc

Previous studies have indicated that Apc inactivation in the intestinal epithelia leading to Wnt pathway-dependent expression of *c-Myc* is the critical mediator of malignant transformation *in vivo* (Dave et al., 2017; Sansom et al., 2006; Sur et al., 2012). We reasoned that blocking c-*Myc* gene activation via Wnt pathway in organoids would attenuate phenotypes imposed by Apc inactivation. We derived organoids from an engineered mouse line lacking a Wnt pathway response element in the *c-Myc* promoter (the *Myc-335*^−/−^ allele; Fig. S4) (Sur et al., 2012). *Myc-335*^−/−^ mice grow normally and importantly, are resistant to intestinal tumorigenesis in an *Apc*^*min*/+^ background (Sur et al., 2012). Wnt3A conditioned media treatment of *Myc-335*^−/−^ and wild-type organoids indicated identical kinetics and frequency of Wnt-oid formation (Fig. 5A, B) that retained the normal Golgi apical localisation (Fig. 5C). Within the 7-day time course of Wnt3A treatment, we observed no changes in growth rate between wild-type and *Myc-335*^−/−^ organoids (Fig. 5D). Taken together, our data indicate that regulation of intracellular organisation and epithelial tissue morphology by Wnt pathway activity is independent of c-Myc expression.

**Figure 5.**
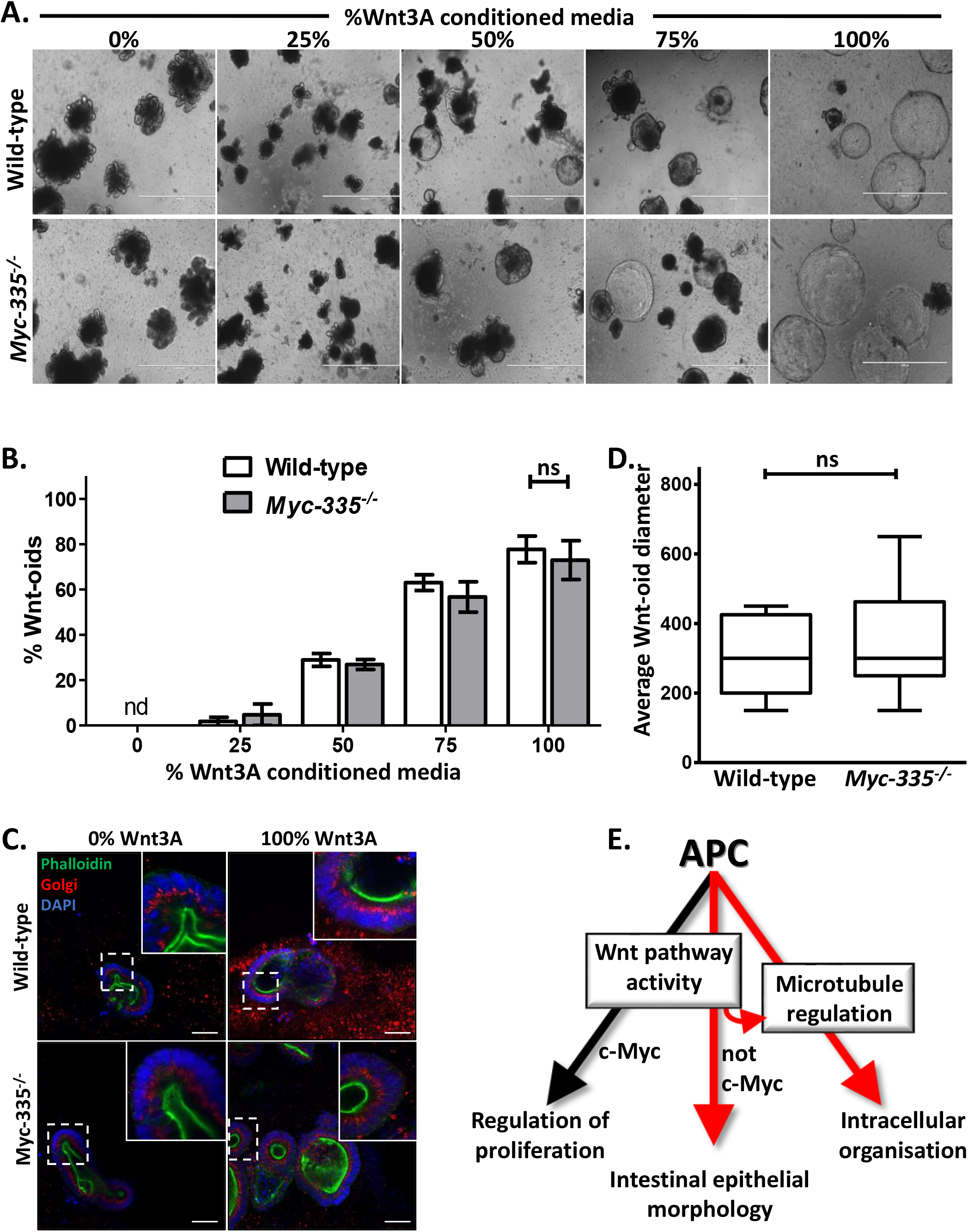
The Wnt pathway target gene *c-Myc* is not a determinant of tissue morphology and intracellular organisation in Apc inactivation. **A.** Representable brightfield images of wild-type and *Myc-335*^−/−^ organoids grown in increasing concentrations of Wnt3A conditioned media for 7 days. Scale bars, 1000 μm. **B.** Quantification of Wnt-oid formation under conditions described above. Data displayed is derived from a minimum of 200 individual organoids from two independent experiments for each concentration of Wnt3A conditioned media. Error bars ± SD; nd – not detected; ns – not significantly different. **C.** Average Wnt-oid diameter (μM) of wild-type and *Myc-335*^−/−^ organoids after 7 days growth in the maximal dose of Wnt3 A conditioned media. Data for the box plots was from greater than 50 organoids from two independent experiments for each organoid type. ns – not significantly different. **D.** Fluorescence confocal microscopy of wild-type and *Myc-335*^−/−^ organoids grown in the absence or presence of the maximal dose of Wnt3A conditioned media for 7 days. All sections were labelled with DAPI (blue), fluorescent phalloidin (*green*) and an antibody to ZFPL1 (red). Scale bars, 50 μm. **E.** Model for Apc regulation of proliferation, tissue morphology and intracellular organisation in the small intestinal epithelia.

## Discussion

In this study, we unmasked individual molecular systems controlled by Apc in the intestinal epithelia through loss-of-function. Oncogenic Apc mutations are the principle driver of colon epithelial tumorigenesis and sufficient for malignant transformation of the colon and small intestinal epithelia. We stratify three emergent phenotypes in the murine intestinal epithelia that are the direct consequence of oncogenic Apc mutations: de-regulated proliferation, disrupted epithelial morphology and compromised microtubule dynamics leading to defective intracellular organisation.

In the intestinal epithelia, Apc activity restricts enterocyte proliferation through stringent control of the Wnt pathway-dependent transcriptional programme. In particular, regulated expression of the Wnt pathway target gene, *c-Myc*, constrains proliferation to discrete, localised niches, providing a key molecular barrier to malignant transformation (Dave et al., 2017; Quyn et al., 2010; Sur et al., 2012); whereas oncogenic Apc mutations in the intestinal epithelia are sufficient to drive neoplastic growth, the absence of *c-Myc* expression attenuates all transforming properties of Apc inactivation, *in vivo* (Sansom et al., 2006). Less well understood is how oncogenic Apc mutations deregulate epithelial morphology and intracellular organisation. We have established that organoids and their Apc-deficient counterparts, tumoroids, are a tractable model that effectively recapitulate the morphological and organisational hallmarks modelling the transition between intestinal epithelia and tumours.

Treatment of organoids with Wnt3A drives their intra-conversion into cystic tumouroid-like structure, termed Wnt-oids that, in contrast to tumoroids, maintain intracellular organisation of the component cells. Our interpretation is that Wnt3A treatment leads to selective inhibition of Wnt pathway regulation by Apc, compromising constraints on epithelial morphology, but retaining the integrity of the microtubule cytoskeleton and intra-cellular organisation – supporting the notion that regulation of epithelial morphology and cytoskeletal integrity are uncoupled. Conversely, selective destabilisation of microtubules compromises intracellular organisation in component organoid cells, yet normal morphology of the epithelia monolayer is retained. Taken together, our data support a model whereby Apc controls enterocyte proliferation and epithelial morphology through Wnt pathway regulation and regulates the microtubule cytoskeleton and intracellular organisation through other, separate pathways (Figure 5D).

How then does Apc regulation of Wnt pathway activity impact the morphology of the epithelial monolayer? Our data support direct control of epithelial morphology by Wnt pathway activity rather than an inability of organisational constraints to cope with exuberant proliferation. In the intestinal epithelia, neoplastic growth is the result of precocious Wnt pathway target gene expression driving deregulated expression of the Wnt pathway target gene, *c-Myc*. Although deregulated *c-Myc* expression is regarded as the major culprit in all transforming phenotypes attributed to Apc loss *in vivo* (Sansom et al., 2007), we find that Wnt pathway-dependent control of *c-Myc* expression has no influence on intestinal epithelial morphology or cellular organisation. Moreover, within the timeframe of our experiments, we did not observe any changes in the rate of proliferation accompanying the intra-conversion of organoids to Wnt-oids. We conclude that compromised epithelial morphology, as a result of Apc inactivation, is not dependent on deregulated *c-Myc* expression, nor is it the result of increased proliferative pressure on organisational constraints on the epithelial monolayer.

It will be important to identify Wnt pathway targets that control intestinal epithelial morphology – we anticipate that targeted modulation of such genes may provide therapeutic value for preventing or even reversing the compromised epithelial morphology accompanying malignant transformation of the intestinal epithelia. The intra-conversion between organoids and Wnt-oids is a ready-made assay system for rapidly testing sufficiency of Wnt pathway candidate target genes by their targeted loss of function; a list of such candidates has been previously identified by Sansom and colleagues (Sansom et al., 2007).

One striking observation was that Apc regulates the integrity of the microtubule cytoskeleton and likely, as a consequence, the intracellular location of organelles such as the nucleus, Golgi, centrosome and intracellular vesicles. Although control of the microtubule cytoskeleton may be mediated directly by the Apc C-terminal microtubule and/or EB1 binding domains (Morrison et al., 1998; Munemitsu et al., 1994), it is also possible that Wnt pathway regulatory components downstream of Apc or even Wnt pathway transcriptional targets contribute to microtubule integrity. For example, truncated Apc in *Apc*^1638T/1638T^ mice retains the ability to regulate Wnt pathway activity and maintain the integrity of the microtubule cytoskeleton (Smits et al., 1999). Our interpretation is that regulation of the Wnt pathway suppresses defects in the microtubule cytoskeleton, *in vivo*. It remains to be determined whether this is the case in the intestinal epithelial-autonomous milieu of *in vitro* organoid culture.

In colon cancer, oncogenic mutations that inactivate Apc are 10-fold more prevalent than oncogenic mutations in other Wnt pathway regulatory components suggesting that functions other than Wnt pathway deregulation contribute to disease aetiology. Although compromised microtubule integrity is a likely consequence of Apc truncations that delete C-terminal microtubule and EB1 binding domains, it is unlikely to impact tumorigenesis—the presence of C-terminal microtubule and EB1 binding domains in truncated versions of Apc has no impact on tumorigenesis (Lewis et al., 2012). However, one intriguing possibility is that compromised microtubule integrity in Apc mutant tumour cells contributes to chromosome instability (CIN). CIN is a feature of the evolution of aggressive colorectal adenocarcinoma right from the outset, evident in the smallest adenomas and multiple reports have directly linked oncogenic APC mutations in CRC with a predisposition to CIN (Dikovskaya et al., 2007; Fodde et al., 2001; Kaplan et al., 2001). Importantly, embryonic stem cells derived from *Apc*^1638T/1638T^ mice develop hallmarks of CIN (Fodde et al., 2001) and overexpression of truncated APC lacking the C-terminal domains in chromosomally stable colorectal cancer cells leads to mitotic defects, including errors in kinetochore attachment and alignment of chromosomes (Green and Kaplan, 2003; Tighe et al., 2004). However, the molecular relationship between Apc loss, microtubule deregulation and chromosome instability in the intestinal epithelia has yet to be established. The experimentally tractable organoid/tumouroid model system we have developed will be invaluable in determining the role of Apc in the loss of microtubule integrity and the impact of CIN in intestinal tumorigenesis.

Our results distinguish individual malignant properties of intracellular disorganisation, compromised tissue morphology and proliferation as direct, but separable consequences of Apc inactivation; we posit that the combination of these emergent properties creates a ‘perfect storm’ for malignant transformation of the rapidly dividing intestinal epithelia, explaining why this tissue is particularly vulnerable to oncogenic Apc mutations.

## Materials and Methods

### Reagents, antibodies and molecular probes

Doxycycline and nocodazole were sourced from Sigma-Aldrich and used at concentrations of 2 μg/ml and 100 nM, respectively. Wnt3A conditioned media was harvested from Wnt3A-expressing L-cells (ATCC, CRL-2647) according to a previously established protocol (Willert et al., 2003). The media was stored for up to two months at 4°C without any detectable loss of Wnt3A activity. Antibodies and molecular probes used for fluorescence microscopy are listed in Table 1.

**Table 1.** Antibodies and molecular probes used for fluorescence confocal microscopy. Except for phalloidin and the antibody to β4-integrin that required the use of fresh-frozen tissue (see below), all fixation prior to labelling with antibodies and molecular probes was performed using 4% formaldehyde.

|  | Conjugate | Company | Catalogue number | Dilution |
| --- | --- | --- | --- | --- |
| **Antibodies** |  |  |  |  |
| Acetylated tubulin | Alexa Fluor 647 | Gift from E. Derivery |  | 1:300 |
| β-catenin | | BD Biosciences | 610153 | 1:200 |
| β-tubulin | | CST | 2146 | 1:200 |
| β4-integrin | | Abcam | ab25254 | 1:200 |
| c-MYC |  | Abcam | ab32072 | 1:1000 |
| Ki67 |  | ThermoFisher | MA5-14520 | 1:200 |
| Lysozyme |  | Dako | A0099 | 1:200 |
| Pericentrin |  | Abcam | ab4448 | 1:200 |
| Phospho-histone 3 (PH3) |  | CST | 9701 | 1:200 |
| Vinculin |  | CST | 4650 | 1:5000 |
| ZFPL1 (Golgi) |  | Sigma | HPA014909 | 1:200 |
| ZO-1 |  | Millipore | MABT11 | 1:200 |
| Rat IgG | Alexa Fluor 488/555 | ThermoFisher | A-11006, A-21434 | 1:500 |
| Rabbit IgG | Alexa Fluor 488/555 | ThermoFisher | A-11008, A-21428 | 1:500 |
| Mouse IgG | Alexa Fluor 488/555 | ThermoFisher | A-21422, A-11001 | 1:500 |
| Rabbit IgG | HRP | Abcam | ab6721 | 1:10000 |
| Mouse IgG | HRP | Abcam | ab6789 | 1:10000 |
| **Molecular probes** |  |  |  |  |
| DAPI |  | SouthernBiotech | 0100-20 |  |
| Phalloidin | Alexa Fluor 488 | ThermoFisher | A12379 | 1:500 |
| UEA-1 (fUEA) | Rhodamine | Vector laboratories | RL-1062 | 1:2000 |

### Tissue preparation and fluorescent labelling

All procedures using mice were performed under the UK Home Office guidelines. Intestines obtained from wild-type CL57BL/6, *Apc*^1638T/1638T^, *Apc*^Min/+^ CL57BL/6, *Apc*^fl/fl^ *LSL tdTo*m (a gift from Winton laboratory) and *Myc-335*^−/−^ (a gift from Taipale laboratory) mice were either fixed in 4% formaldehyde and embedded in paraffin or fixed-frozen in 10% formalin and embedded in optimal cutting temperature (OCT) liquid, followed by snap freezing (fresh-frozen tissue).

Small intestinal epithelial sections (4% formaldehyde-fixed or fresh frozen) for molecular probe and antibody labelling were cut at a thickness of 4.5 μm onto slides. The exception was slides labelled with β-tubulin or acetylated-tubulin in which case 20 μm thick formaldehyde-fixed sections were cut onto poly-L-lysine coated slides.

For formaldehyde-fixed samples, epitope retrieval was performed in sodium citrate buffer (sodium citrate 10 mM, 0.05 % Tween-20, pH 6.0). Primary antibody incubations were carried out at 4°C overnight and secondary antibody incubation for 2 hours at room temperature, both in PBS containing normal goat serum (5%) and 0.1% Tween20. Samples were mounted in DAPI-containing Fluoromount-G (Thermo-Fisher).

### Organoid preparation and fluorescent labelling

Murine small intestinal epithelial organoids were derived from the ileum of mouse small intestine according to Sato et al. 2013 (Sato and Clevers, 2013). Tumoroids were derived from tumours within the ileum of 110 day-old *Apc*^Min/+^ mice (Haigis et al., 2004). All organoids and tumoroids were cultured according to Urbischek et al. 2018 (Urbischek et al., 2019).

Organoids were seeded in Matrigel onto eight-well chamber slides (ThermoFisher) 48 hours prior to fluorescent labelling. Organoids were fixed in 92% methanol containing 8% formaldehyde and labelled following a published protocol (Goldspink et al., 2017). Organoids in primary antibody were incubated at 4°C overnight. The next day the slides were incubated at room temperature for 1 hour (allowing the Matrigel to harden), washed and then incubated for 1 hour at room temperature in the secondary antibody. Labelled organoid samples were then mounted in DAPI-containing Fluoromount-G.

### Organoid and tissue imaging and data analysis

Fluorescent imaging of tissue was carried out using a Nikon C2 plus confocal microscope using 40X objective lens. Images were processed using ImageJ software. Fluorescent labelling of each antibody was repeated a minimum of three times.

Imaging of organoids was done using Nikon C2 plus confocal microscope using the 20X and 40X objectives and an automated spinning disc confocal microscope (YOKOGAWA Cell Voyager CV8000) using 40X objective. Z-stacks were taken at 1 μm steps. Images were processed and published using ImageJ software. All figures presented are representative images from a single plane within the Z-stack of the imaged specimen. For the quantification of organelle positioning within organoids, approximately 200 cells were counted per experiment manually.

### Plasmids and organoid expression

The *piggybac* transposon and *tet-on* expression system was a kind gift from Bon-Kyoung Koo. The previously validated shRNA targeting mouse *Apc* (Dow et al., 2015) was inserted into the tet-responsive shRNA expression vector, pB-TRE-IRES-mCherry. The three plasmid system also consists of pB-CAG-rtTA, the vector for constitutive rtTA expression, and pPiggybac, the expression vector for constitutive expression of the *piggybac* transposase (Fujii et al., 2015).

The shApc organoid line was generated by transfection of the pB-TRE-shApc-IRES-mCherry, pPiggybac and pB-CAG-rtTA plasmids (Fig. S3A) using a NEPA21 electroporator according to a previously published protocol (Fujii et al., 2015). Organoids were selected for integration of constructs in organoid media containing Wnt3A conditioned media (Urbischek et al., 2019) supplemented with 150 μg/ml Hygromycin B (ThermoFisher) for 7 days, after which the media was switched to organoid media (Urbischek et al., 2019).

### Validation of shApc organoid line

For western blotting, organoids were recovered from Matrigel using several rinses of ice-cold phosphate buffered saline (PBS) and the pellet was lysed with 50 μl 1X RIPA buffer (Millipore) containing protease (Sigma) and phosphatase inhibitors (Roche). Samples were loaded onto NuPAGE 3-8 % Tris-Acetate gradient gels (ThermoFisher) prior to transfer onto PVDF membrane. Antibodies used for probing membranes in PBS containing 0.2% Tween 20 and 5% non-fat milk are in Table 1.

Expression levels of Apc and mCherry were determined by qRT-PCR. RNA was isolated from organoids and tumoroids using the ReliaPrep RNA Cell Miniprep System kit (Promega) and cDNA was prepared using the High Capacity cDNA Reverse Transcription kit (ThermoFisher) all according to manufacturers’ instructions. qRT-PCR was carried out using Fast SYBR Green Master Mix using a QuantStudio 5 real-time PCR sytem (both Applied Biosystems). B2m was used as a housekeeping gene and relative fold changes in Apc and mCherry expression were derived from ∆∆CT. The following primers were used: Apc (Forward: AGCCATGCCAACAAAGTCATCACG; reverse: TTCCTTGCCACAGGTGGAGGTAAT), mCherry (Forward: CACGAGTTCGAGATCGAGGG; reverse: CAAGTAGTCGGGGATGTCGG) and B2m (Forward: ACCCCCACTGAGACTGATAC; reverse: ATCTTCAGAGCATCATGATG).

## Acknowledgements

The authors thank Nami O. Yamada and Wenduerma for technical assistance, Thomas Foets and Manuela Urbischek for discussions of the work and Trevor Littlewood for critical review of the manuscript. This work was supported by a PhD studentship project funded by AstraZeneca to H.R.

## Supplemental Figures

**Supplementary Figure 1.**
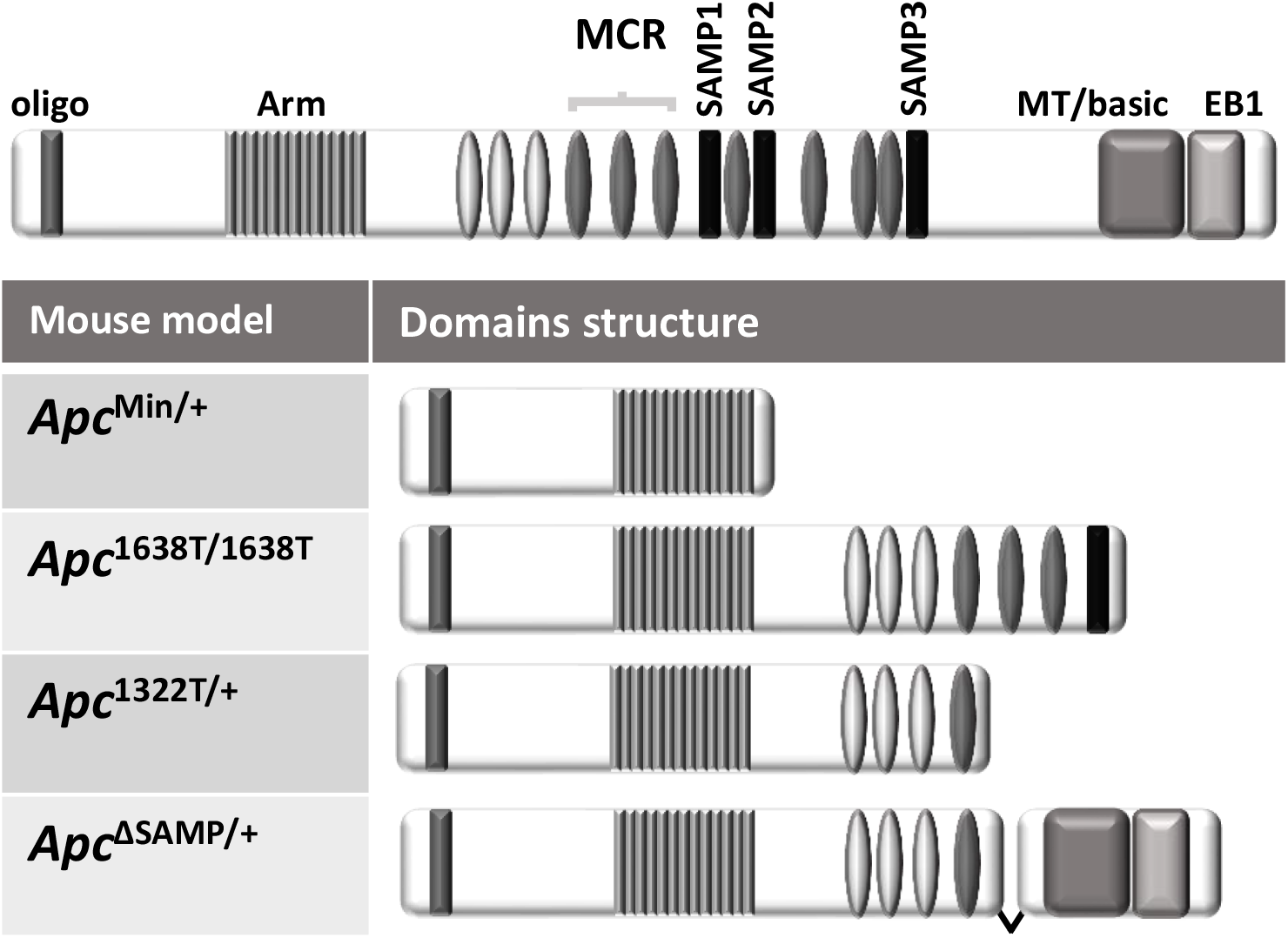
Domain structure of Apc and mutant variants expressed in mouse models of intestinal epithelial tumorigenesis. Labels for Apc protein-interaction domains are as in Fig. 1C.

**Supplementary Figure 2.**
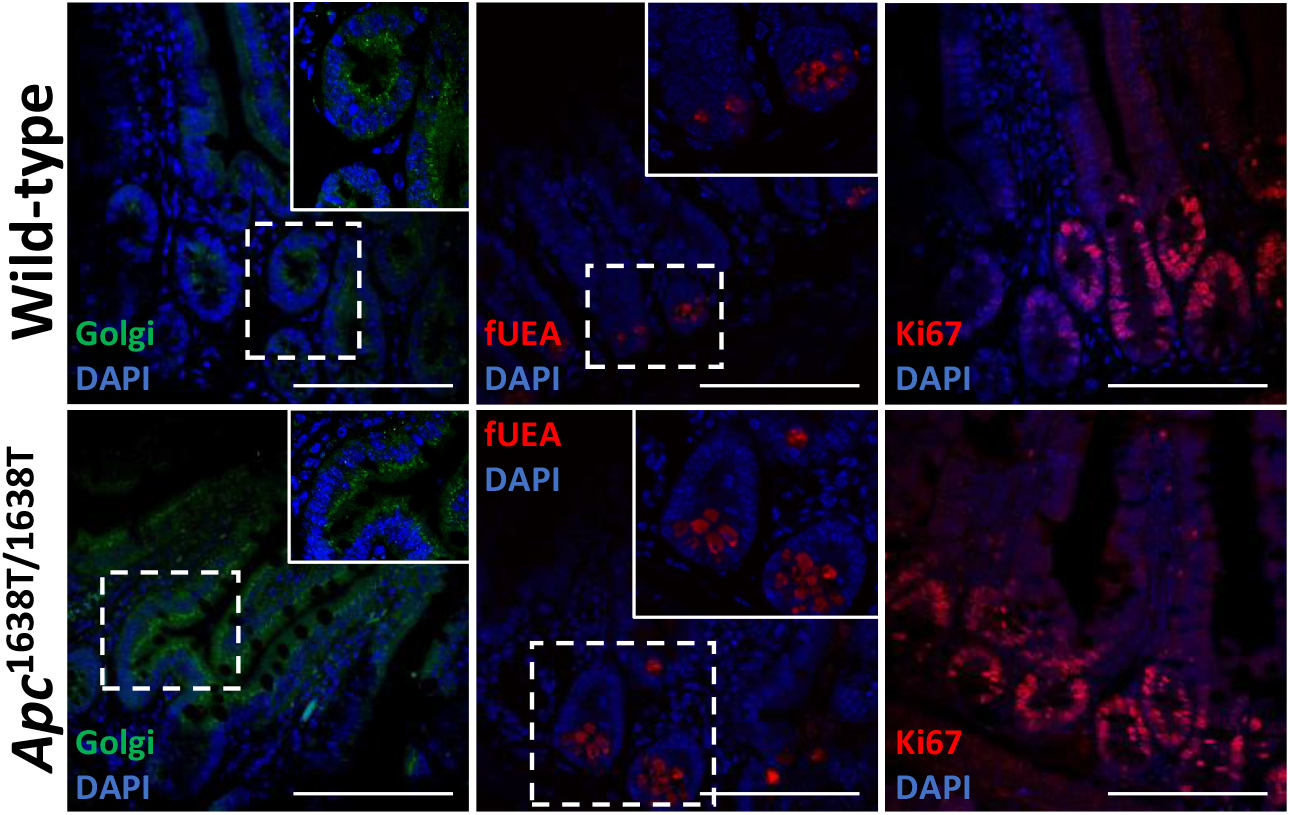
No loss of microtubule organisation and intestinal epithelial morphology in *Apc*^1638T/1638T^ mice. Fluorescence confocal microscopy of sections of small intestinal epithelia from a wild-type (top panels) and *Apc*^1638T/1638T^ (bottoms panels) mouse. *Left panels* are labelled with an antibody to ZFLP1 (green); *middle panels* are labelled with fUEA (red); right panels are labelled with an antibody to Ki67 (red). All sections were co-labelled with DAPI. Scale bars, 100 μm.

**Supplementary Figure 3.**
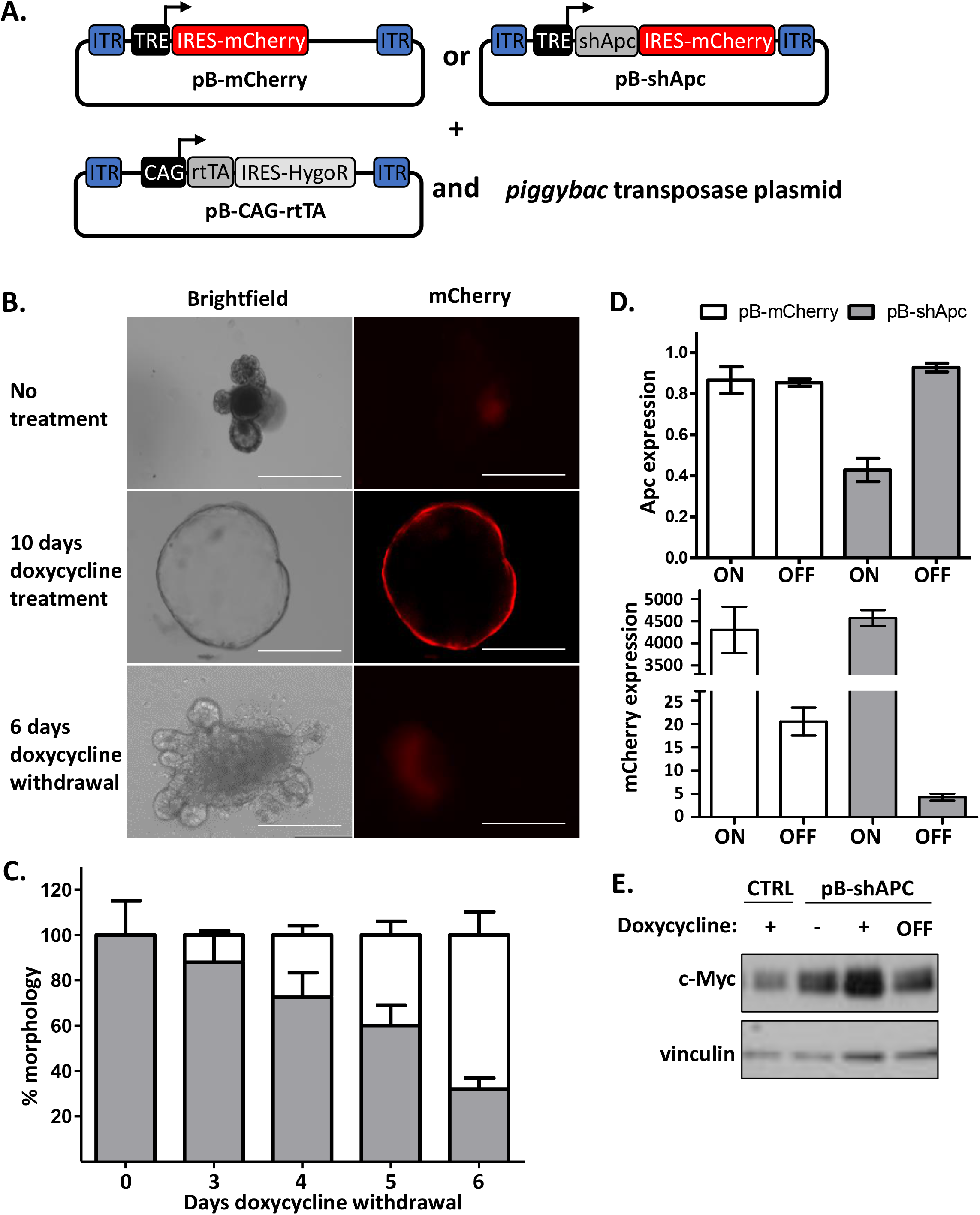
The pB-shApc switchable model of *in vitro* tumorigenesis/tumour regression recapitulates the phenotypic consequences of oncogenic Apc mutations. **A.** Transgenes used for the construction of the control pB-mCherry or pB-shApc organoid lines. The shApc expression system includes pB-CAG-rtTA for constitutive rtTA expression and expression of the *piggybac* transposase for stable integration into organoids. In-built *tet-on* system enables inducible expression of shApc linked to mCherry by treatment of pB-shApc organoids with doxycycline. **B.** Time-course of doxycycline treatment of pB-shApc organoids; after ten days all organoids have converted to spheroids accompanied by mCherry expression. Subsequent doxycycline withdrawal and growth for an additional 6 days leads to restoration of organoid morphology. Scale bar, 200 μm. **C.** Quantification of spheroid conversion to organoids upon doxycycline withdrawal. Data is the average of greater that 100 organoids from two independent experiments were scored for each timepoint. **D.** Control or pB-shApc organoids were treated with doxycycline as above and protein lysates were probed with antibodies to c-Myc and the loading control vinculin. **E.** QRT-PCR quantification of Apc and mCherry expression in control or pB-shApc organoids after two days doxycycline treatment, ‘ON’, or 6-days post withdrawal, ‘OFF’. Data is represented as mean of two independent experiments. Error bars ± SD

**Supplementary Figure 3.**
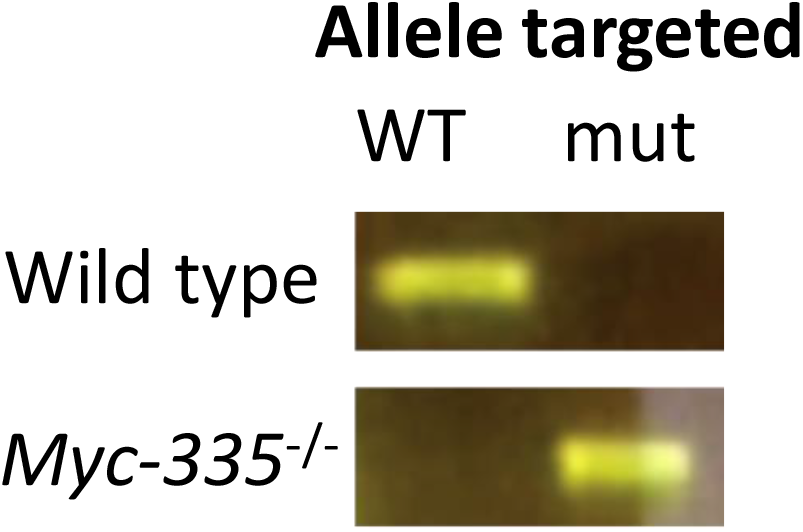
Genotyping of intestinal epithelia from *Myc-335*^−/−^ mouse used in this study according to Sur et al. PCR genotyping of *Myc-335*^−/−^ intestinal epithelia using primers targeting the wild-type (WT) or *Myc-335* mutant allele (mut). Primers and genotyping conditions are described in Sur et al. (Sur *et al.*, 2012).

